# μCas, a novel class of miniature type-V Cas12f nucleases with diverse PAM

**DOI:** 10.1101/2023.02.03.527072

**Authors:** Allison Sharrar, Luisa Arake de Tacca, Trevor Collingwood, Zuriah Meacham, David Rabuka, Johanna Staples-Ager, Michael Schelle

**Author notes:** Allison Sharrar and Luisa Arake de Tacca contributed equally to this work. Corresponding Author: Michael Schelle.

## Abstract

Small CRISPR-Cas effectors are key to developing gene editing therapies due to the packaging constraints in viral vectors. While Cas9 and Cas12a CRISPR-Cas effectors have advanced into select clinical applications, their size is prohibitive for efficient delivery of both nuclease and guide RNA in a single viral vector. Type-V Cas12f effectors present a solution given their small size. Here we describe μCas, a novel class of miniature (<490AA) type-V Cas12f nucleases that cleave double stranded DNA in human cells. We determined their optimal trans-activating RNA (tracrRNA) empirically through rational modifications which resulted in an optimal single guide RNA (sgRNA). We show that the μCas nucleases have broader PAM preferences than previously known Cas12f effectors. The unique characteristics of these novel nucleases add to the diversity of the miniature CRISPR-Cas toolbox while the expanded PAM allows for the editing of genomic locations that could not be accessed with existing Cas12f nucleases.

## Introduction

Clustered regularly interspaced short palindromic repeat (CRISPR) associated (Cas) systems (CRISPR-Cas) are powerful tools to edit DNA and RNA in a targeted manner.^1–4^ CRISPR-Cas, which functions as adaptive immunity for bacteria and archaea in nature, uses diverse Cas-protein effectors and gene organizations.^5–7^ New CRISPR-Cas systems, identified through extensive metagenomic efforts, have added to the understanding of how they aid bacteria and archaea against invading viral threats.

There are two classes of CRISPR systems with multiple CRISPR types. The first system to be used for mammalian editing was the class II, type II CRISPR-Cas9 system.^1,2,7^ Class II, type V-A systems, including the Cas12a nucleases, were discovered in 2016.^8^ The initial discovery included several examples of active effectors such as AsCas12a from *Acidaminococcus* sp., LbCas12a from *Lachnospiraceae bacterium* and FnCas12a from *Francisella novicida*. Both Cas9 and Cas12a nucleases are capable of cleaving doublestranded DNA, leading to mutations after DNA repair. However their large size (ranging from 120 kDa to 160 kDa) prevents optimal packaging into a single adeno-associated viral particle (AAV), limiting their use for *in vivo* therapeutics.

Recently, small CRISPR-associated effector proteins belonging to the type V-F subtype (Cas12f) were identified through the mining of sequence databases.^7,9–11^ The first Cas12f (Un1Cas12f1, also known as Cas14a) is referred to as a miniature CRISPR-Cas due to its small size (60 kDa) compared to Cas9 and Cas12a proteins.^9^ Recent classifications have grouped this family into Cas12f1 (Cas14a and type V-U3), Cas12f2 (Cas14b) and Cas12f3 (Cas14c, type V-U2 and U4).^12^ To date, only three Cas12f effectors have been shown to edit mammalian systems, Un1Cas12f1, AsCas12f, and SpaCas12f1. Cas12f enzymes require a tracrRNA sequence as well as a crRNA in their guide structures. They cleave double stranded DNA and have a 5’ T-rich PAM preference, leaving an overhang at the cut site.^13–15^

In this work we describe the discovery of μCas, a novel class of miniature Cas12f nucleases most closely related to Cas12f1 with robust editing in human cells. These novel enzymes recognize a broader PAM motif than Un1Cas12f1, increasing the density of genome targeting sites. μCas nucleases expand the potential of miniature CRISPR-Cas genome editing to treat human diseases.

## Materials and methods

### Nuclease discovery

Candidate nucleases were discovered by searching Cas12f sequences known to edit in mammalian cells^13,15,16^ against the NCBI non-redundant (nr) and metagenomic (env_nr) protein databases using BLAST.^17^ The genomic source sequences of the nucleases were then searched for CRISPR arrays and other Cas genes. CRISPR arrays were identified with CRISPRDetect^18^ and a local version of CRISPRCasTyper.^19^ Known Cas genes were searched for using hidden Markov model (HMM) profiles built from Pfam^20^ seed sequences with the HMMer suite.^21^

### Cas12 phylogenetic tree

Cas12 sequences from subtypes a-k were collected from source literature^5,8,9,22–25^ and aligned with Clustal Omega v1.2.4.^26^ A phylogenetic tree was built off the alignment using RAxML v8.2.12 with the PROTGAMMALG model and 100 bootstrap samplings^27^ The tree was visualized and edited with iTOL v6.6.^28^

### tracrRNA Design

TracrRNA sequences are known to bind to the direct repeat and lie between the end of the nuclease and start of the CRISPR array.^13,16^ Therefore, tracrRNAs specific to our nucleases were estimated from this region of their genomic source sequences. The 3’ end of the tracrRNA was estimated from where a portion of the direct repeat binds to this region, as determined by nucleotide BLAST.^17^ The 5’ end of the tracrRNA was estimated based on total lengths and sequence motifs (like 5’-NTTC) of known Cas12f tracrRNAs. To make the sgRNA, the tracrRNA is connected to the binding portion of the direct repeat by an artificial loop. Folding patterns of the sgRNA designs were then predicted with the UNAFold web server.^29^ Based on predicted folds, the 5’ end of the tracrRNA was sometimes revised to create completed stem loops. While preserving the structure, single T’s were replaced with A’s to disrupt Poly-T regions throughout the sgRNA.

### Plasmid construction

Codon-optimized genes encoding candidate nucleases were synthesized and cloned into the mammalian expression vector under the CMV promoter, pTwist_CMV (Twist Biosciences). The cloned nucleases were placed into the expression vector with a SV40 Nuclear Localization Sequence (NLS) fused to the N-terminus and a nucleoplasmin NLS on their C-terminus, followed by a 3x HA tag. A similar vector was created with Un1Cas12f1. To make specific sgRNA vectors for each nuclease, direct repeat sequences were placed downstream of the matching tracrRNAs and the combination of tracrRNA-Direct repeat-Spacer was placed downstream of a U6 promoter with a starting G.; Several different tracrRNA and Spacer configurations were tested for each nuclease and their sequences are shared on the supplemental materials.

### Editing activity in human cells

Nucleases were tested for activity in HEK293T cells following plasmid transfection using Mirus Transit X2 reagent. Tests were performed in 96 well plates transfected with 100 ng of nuclease expression vector and 100 ng of targeting guide vector following the Mirus Transit X2 transfection recommendations. Samples were incubated for 72h and harvested with Quick Extract (Lucigen). Genomic DNA was amplified using genomic region-specific primers (Supplemental). Samples were checked on a 2% agarose gel for purity, cleaned up and sequenced by Sanger sequencing. TIDE analysis was performed following the published method^30^ and performed according to recommendations at tide.nki.nl. TIDE output data on editing efficiency was plotted using Prism software.

### Off-Target Editing Activity

The nuclease μCas-116 was tested with either a guide matching the TCRA gene or a guide with a single mismatch for TCRA at different positions. The mismatched guides act as artificial off-targets to determine the propensity of the nuclease to edit with mismatches at each position of the guide. Editing efficiency was measured for the matched guide and mismatched guides by Sanger sequencing followed by TIDE analysis as described previously. Non-transfected (NT) cells were also harvested, amplified, and sequenced via the same methods to set a limit of detection (L.O.D.), under which editing levels cannot be determined.

### PAM preference determination

To determine the PAM preference of our nucleases, we used the protocol described by Walton and collaborators in 2021.^31^ We used the spacer listed in the article as “Spacer 3” with 10xN in the sequence. We adapted the protocol to use a puc19 vector from NEB instead of p11-LacY-wtx-1 and performed linearization with BsaI. Cell lysis was performed with Buffer NP40 (50mM Hepes pH 7.5, 150mM KCl, 2mM EDTA, 0.5% NP40, 1mM DTT, and 1 Roche protease inhibitor tablet for 10 mL of buffer).

## Results

### Discovery of μCas effectors

Three novel Cas12f effectors, dubbed μCas due to their small size, were discovered in metagenome-assembled genomes (MAGs) from ruminant microbiomes (NCBI BioProject PRJNA657473). Two of the effectors are from the genus *Alistipes* and share 85% protein sequence identity (μCas-116 and μCas-117). μCas-122 is from the genus *Tidjanibacter* and shares 80% protein sequence identity to the others. They are smaller than most known effectors in the Cas12 family at 486-488 aa. Sequence alignment shows a RuvC nuclease domain near the C-terminus, but no greater than 22% protein identity to known Cas12f effectors (Fig. 1).^9^ Their CRISPR arrays contain 29-32 bp spacers and 36 bp repeats sharing a minimum of 66% nucleotide identity with a consistent ATATCCAAC-3’ motif. A putative tracrRNA lies between the nuclease and CRISPR array and there are no other known Cas genes present in the operon (Fig. 2a).

**FIG. 1.**
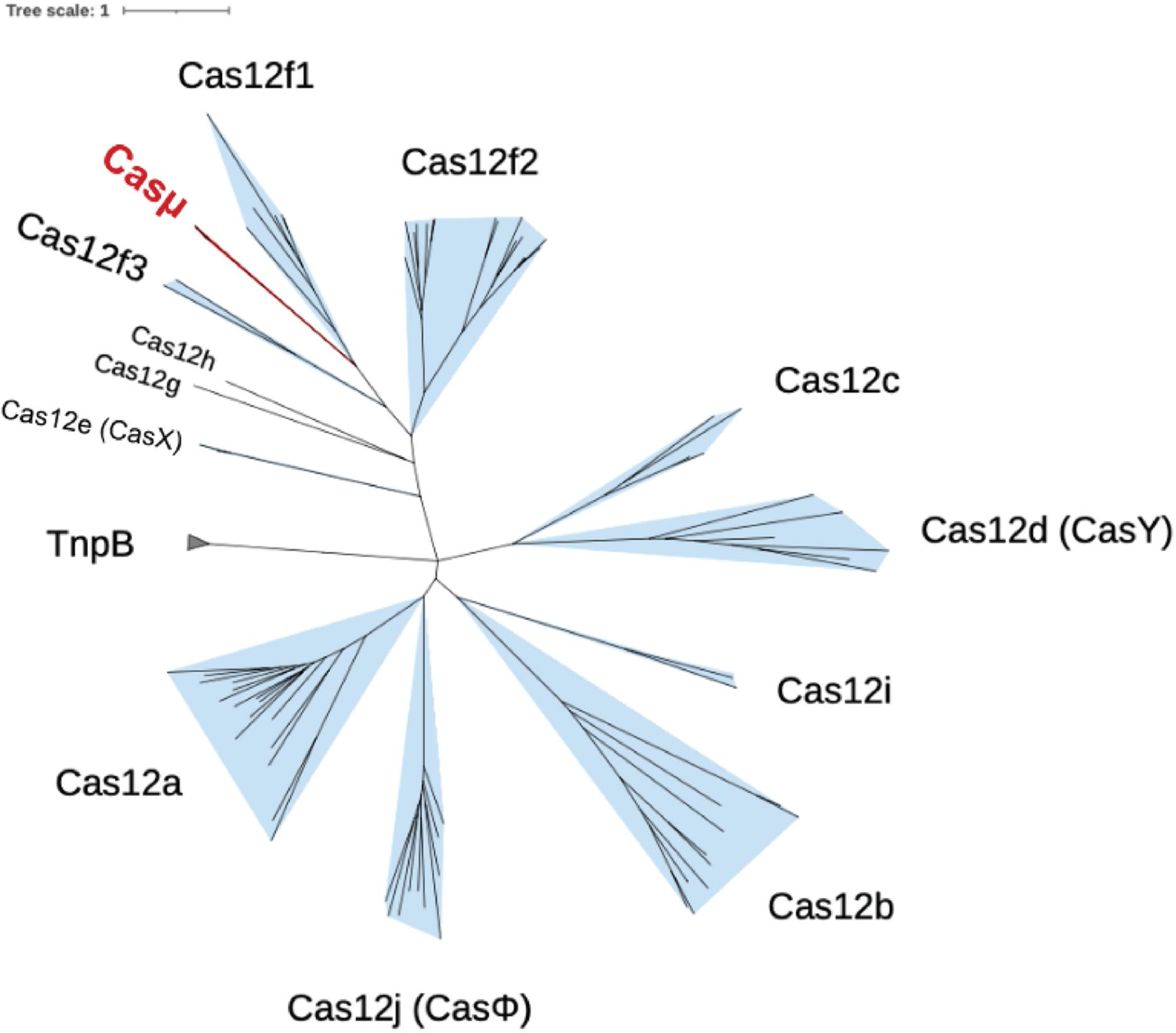
Maximum likelihood phylogenetic tree of known Cas12 subtypes a-j with newly discovered μCas nucleases (μCas116, μCas-117, and μCas-122) in red.

**FIG. 2.**
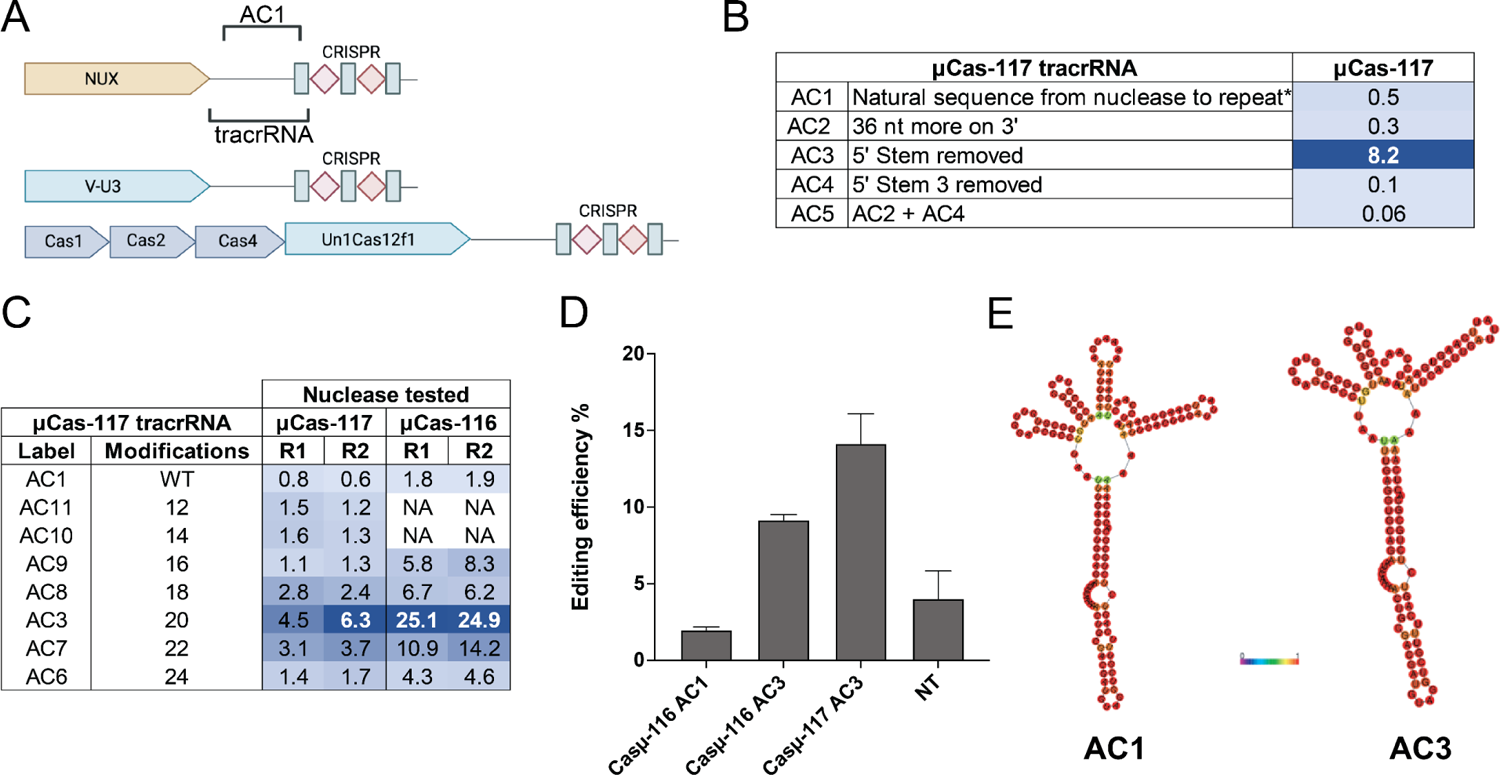
(**A**) Schematic representation of CRISPR-Cas loci and effector protein for type V systems exemplified by V-F (Un1Cas12f1) and V-U3 system. (**B**) Change in percent editing of nuclease based on sgRNA modifications of the tracrRNA. (**C**) Fine truncations on the tracrRNA of the sgRNA with different editing activities with two closely related nucleases. (**D**) Editing efficiency of nuclease μCas-116 with different tracrRNA configurations. (**E**) Difference in tracrRNA prediction of the structure of the native tracrRNA AC1 and the engineered tracrRNA AC3.

### Optimal tracrRNA improves editing in mammalian cells

To determine the optimal tracrRNA needed for editing in mammalian cells, we started with the native genomic location downstream of the nuclease μCas-117, here called AC1. This location has been described to contain the tracrRNA sequence of other Cas12f nucleases (Fig. 2A). We tested different tracrRNA configurations following previous work from Kim and collaborators as well as Wu and collaborators^13,14^ to find a preferred composition for editing in human cells (Fig. 2B). After determining that the version AC3, which has a truncated 5’ stem, was better for editing (Fig. 2B), we made fine truncations in that construct to gain more insight into the RNA space around that tracrRNA (Fig. 2C). These constructs were tested with a TTTG PAM spacer which was previously described to work with UnCas12f.^16^ We also assessed μCas-116, another μCas effector closely related to μCas-117. When tested with its own natural tracrRNA genomic location similar to AC1 for μCas-117, μCas-116 showed minimal editing (Fig. 2D). However, when tested with the μCas-117 AC3 tracrRNA, μCas-116 showed improved editing efficiency (Fig. 2D). Removing the 5’ stem on its natural tracrRNA did not outperform μCas-117 AC3 (Fig. 2D). Interestingly, the overall fold and base-pairing of AC1 and AC3 are similar with most of the sequence showing high base-pairing probability due to canonical interactions on the stems. The main structural difference between the native and engineered tracrRNA is an extra loop on the 5’ of AC1 (Fig. 2E).

With the goal of understanding the most important regions of the tracrRNA portion of the sgRNA, we made a variety of truncations on the main stem. Here we call the truncated pieces “TCa” on the 5’ side of the stem, “TCb” on the 3’ of the stem or “TCab” which has both truncated (Fig. 3A). We observed that all truncations disrupt editing to a certain level, with the most disruptive being regions 1 and 2 on the base of the stem (Fig. 3A and 3B).

**FIG. 3.**
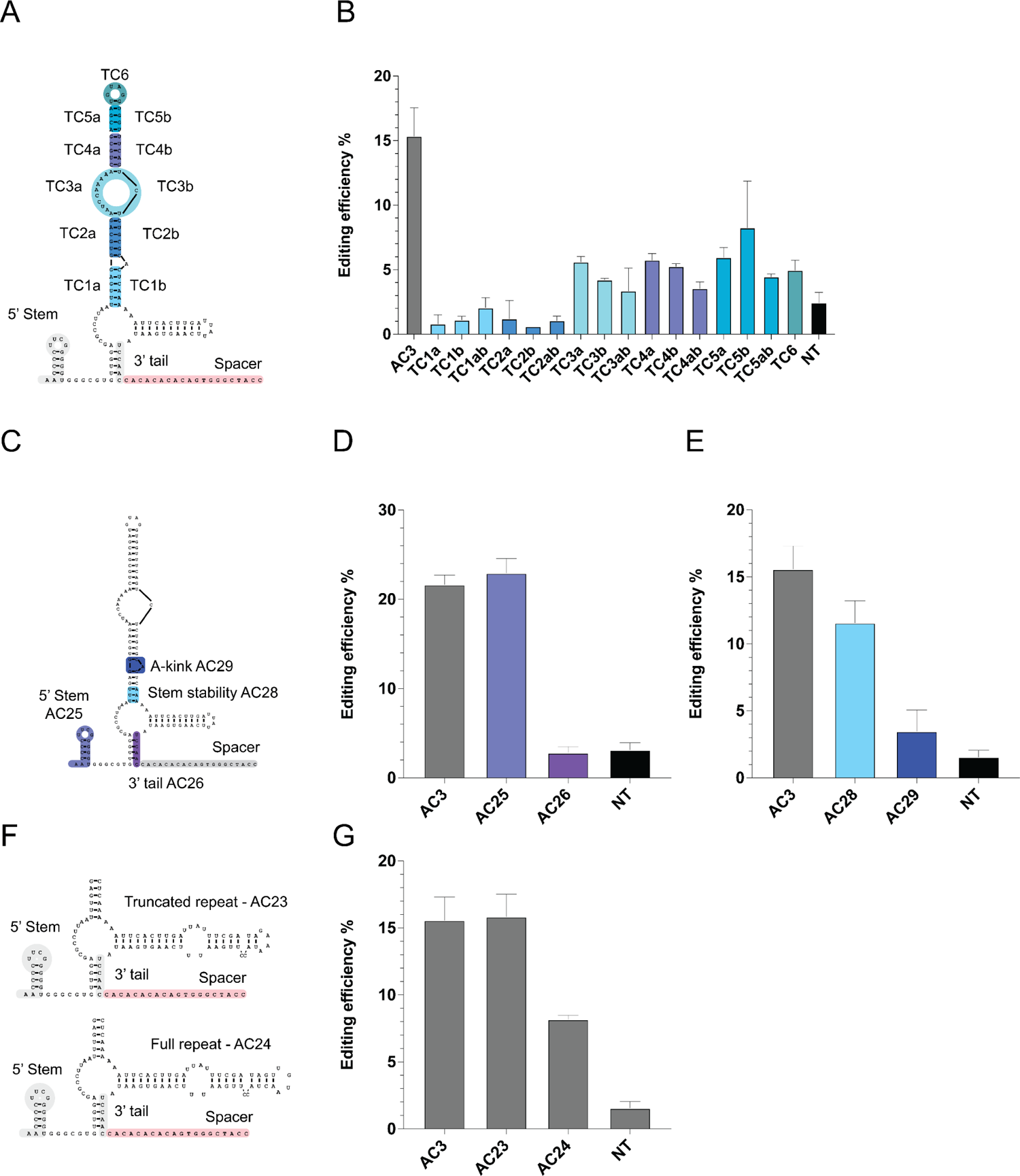
TracrRNA modifications. (**A**) Truncations of middle regions of the third and main RNA stem of tracrRNA AC3. (**B**) Editing efficiency of tracrRNA truncations shown in panel A using μCas-116. (**C**) Graphical representation of the tracrRNA modifications to strengthen the stability of stem. (**D**) Editing efficiency of tracrRNAs shown in panel C. (**E**) Editing efficiency of modifications to strengthen the stability of the tracrRNA shown in panel C using μCas-116 and μCas-122. (**F**) Graphical representation of the different lengths of repeat used to fine tune human cell editing of μCas-116 with AC3-like tracrRNA. (**G**) Editing efficiency of tracrRNA AC3 with different lengths of repeat such as shown in panel F using μCas-116. All editing efficiencies shown in panels B, D, E and G were calculated with the use of TIDE software and sequenced with Sanger methodology.

We initially designed the 5’ start of the tracrRNA based on similar sequences across distinct Cas12f nucleases. To investigate the importance of the conservation at this location, we truncated the stem on the 5’ side of the tracrRNA, making an overall shorter tracrRNA (AC25). This modification slightly improves editing for both μCas-116 and μCas-122 (Fig. 3C and 3D). We then tried to strengthen the base-pairing of the tracrRNA given the importance of the base of the stem observed in Fig. 3A. We modified the A-T base pairs into G-C pairs in the base of the stem shown in green “Stem stability” (AC28) (Fig. 3C). We also removed the kink inserted by an unpaired single A nucleotide (AC29) (Fig. 3C). Improving stability of the stem changed the predicted ΔG of the structure, hinting to a more stable fold but it did not improve the editing efficiency of the nuclease. Removing the A-kink completely abrogated editing capabilities of the nuclease, suggesting that the kink is an important structural feature of the tracrRNA (Fig. 3D and 3E).

We then investigated the importance of the length of the direct repeat for tracrRNA efficiency. We made two alternative configurations in which we used the optimal version with the 5’ stem removed, such as in AC25, and tested a short version of the repeat (AC23) or the full length of the repeat (AC24). We observed that having the full direct repeat was detrimental to editing activity when compared to the truncated version of the repeat (AC3 or AC23) and (Fig. 3C).

After testing different sgRNA configurations, we observed that the μCas-117 tracrRNA AC3 worked optimally with all three μCas nucleases. This sgRNA can be further shortened with a 5’ stem truncation (AC25) without any loss in activity.

### Mismatch tolerance and optimal spacer length

To assess the fidelity of our novel μCas nucleases, we tested the tolerance to mismatches between the guide and target DNA in human cells. We designed a synthetic mismatch panel, in which a single base pair mismatch was created at every position of the TCRA spacer to mimic a single mismatch off-target site (Fig. 4A). We observed that μCas-116 does not tolerate mismatches in nucleotides 1-7 of the spacer (Fig. 4B). Further, μCas-116 shows higher fidelity than Un1Cas12f1, which edits mismatches at positions 1-3 and position 5.^16^ These data suggest that μCas is more stringent than Un1Cas12f1 in the seed region of the spacer.

**FIG. 4.**
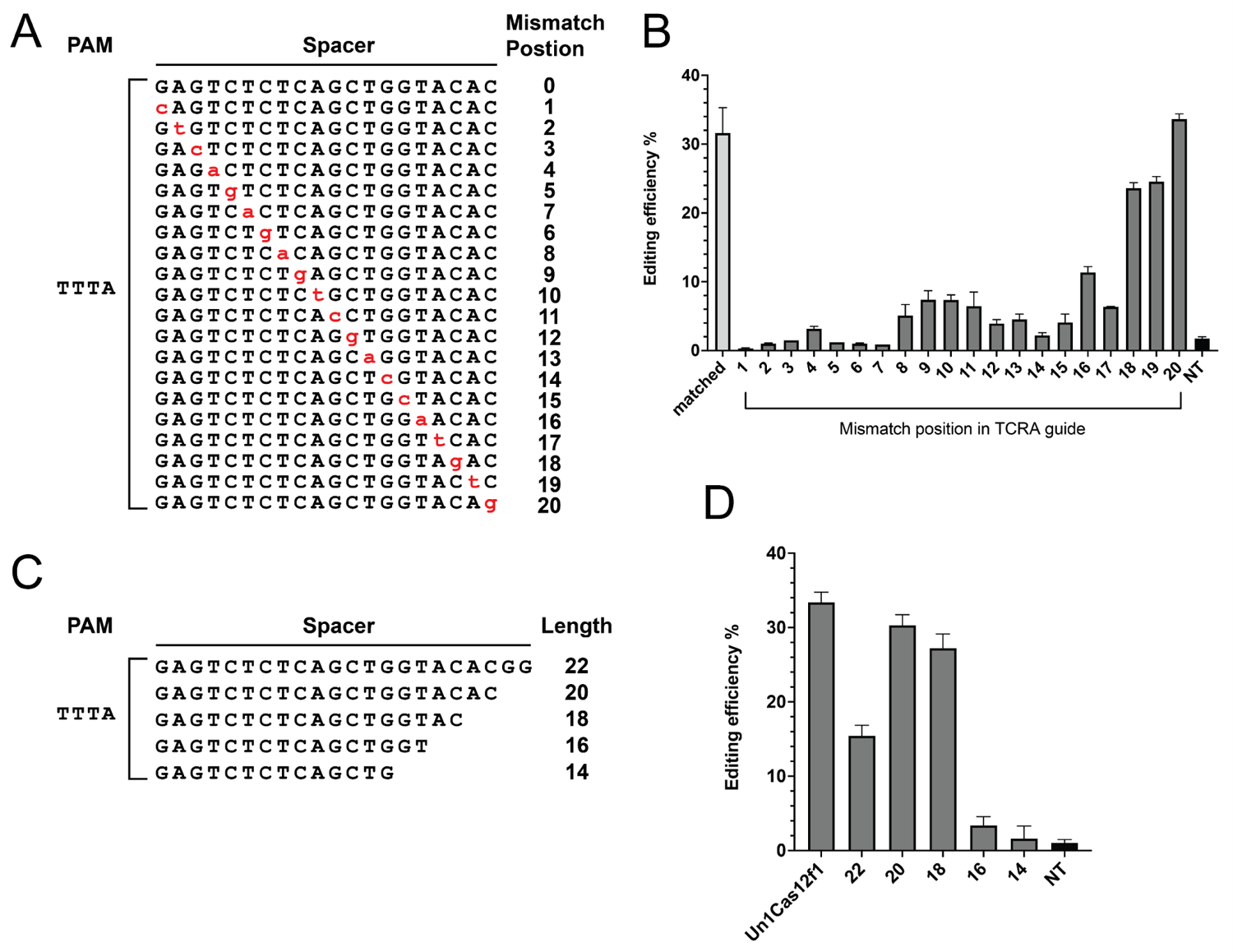
Characterization of μCas nucleases. (**A**) Graphical representation of the mismatches inserted into the spacer to mimic an off-target panel using the TCRA spacer and edited by μCas-116. (**B**) Editing efficiency of spacers with mismatches in every position, as well as the matched spacer for μCas-116. (**C**) Graphical representation of the different lengths of the TCRA spacer to determine the preference of μCas-116. (**D**) Editing efficiency of the different length of spacers with μCas-116. Un1Cas12f1 is used as a positive control and NT stands for non-targeted cells.

We also verified the length of the spacer that our nuclease preferred by truncating the spacer on the PAM-distal side from 22 nucleotides to 14 nucleotides (Fig. 4C). Like other members of the Class II CRISPR system, our novel μCas nucleases prefer spacers of 19-20 nucleotides in length and completely lose activity with spacers that have 16 or fewer nucleotides (Fig. 4D).

### PAM determination and validation of preferences

To learn more about the protospacer adjacent motif (PAM) requirements for DNA targeting of our μCas nucleases, we performed the PAM depletion assay as described by Walton and collaborators.^31^ Briefly, a spacer targeting a randomized PAM plasmid library was incorporated downstream of the tracrRNA and repeat regions of the sgRNA. The preferred PAM sequences for our nucleases were depleted *in vitro*, and the non-preferred PAM sequences were identified by next-generation sequencing (NGS). Interestingly, the μCas nucleases have a distinct PAM preference of DTTR (D = A, G, or T; R = A or G) with a stronger bias towards ATTA PAMs (Fig. 5A). This result differentiates the μCas nucleases from the previously described Cas12f family members Un1Cas12f1, AsCas12f and SpaCas12f1. Un1Cas12f1 and AsCas12f have a PAM preference that is limited to TTTR (Fig. 5A) while SpaCas12f1 prefers NTTY PAM sequences. The PAM preference for μCas nucleases was validated by measuring the editing efficiency at multiple genomic locations with diverse PAM sequences (Fig. 5B). The μCas nucleases are uniquely able to edit a variety of PAM sequences that are inaccessible to Un1Cas12f1.

**FIG. 5.**
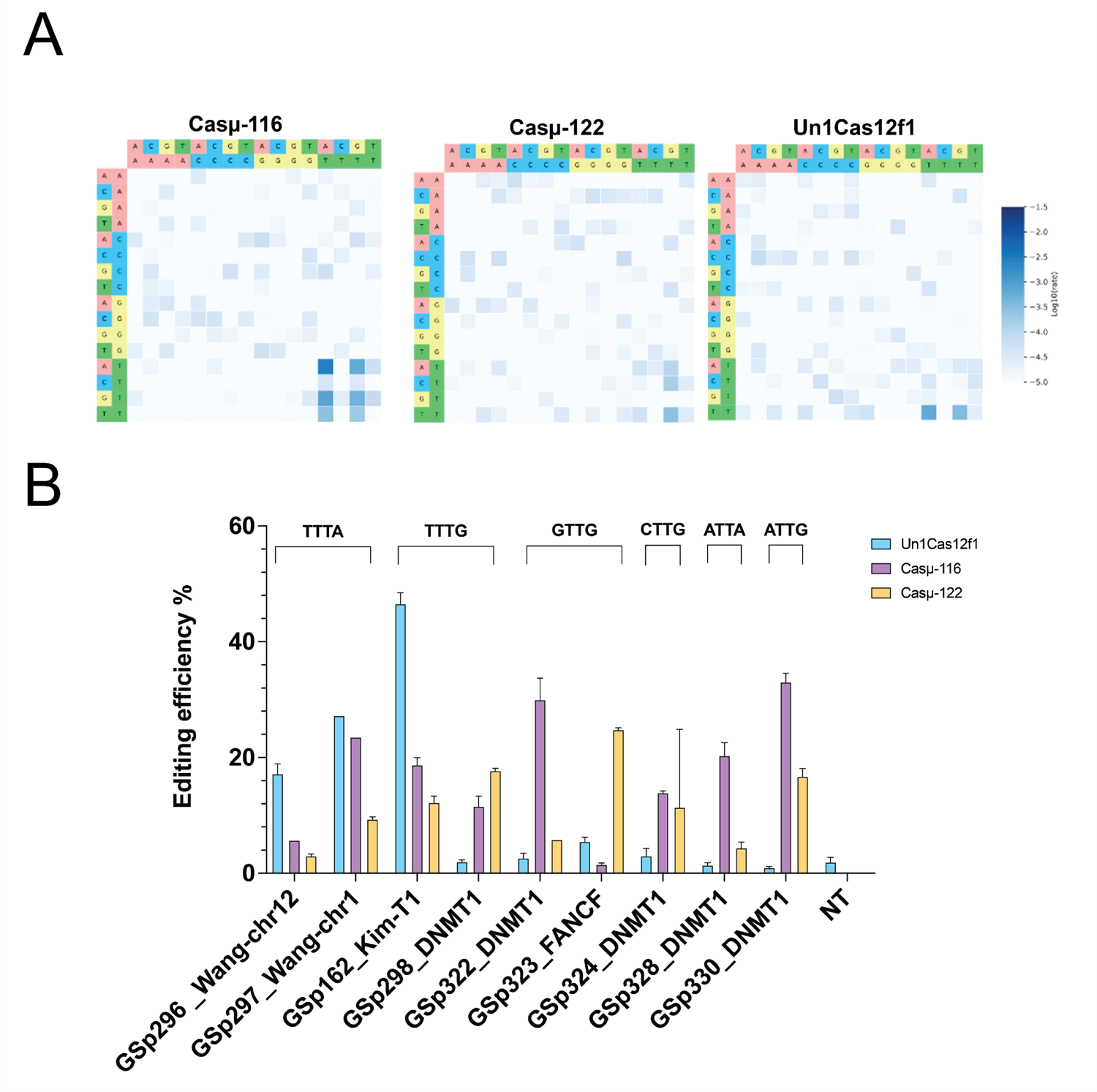
PAM preference and validation. (**A**) PAMDA characterization of μCas-116 and μCas-122 as well as Un1Cas12f1. (**B**) Editing confirmation of PAM preferences of μCas-116, μCas-122 and Un1Cas12f1.

## Discussion

In the present work, we discovered and characterized three miniature type-V Cas12f nucleases which we termed μCas. μCas nucleases are compact editors that are active in human cells and work optimally with a common sgRNA. These nucleases belong to a different clade than Cas12f1, Cas12f2 or Cas12f3 effectors and possess distinct PAM preferences relative to other Cas12f effectors known to edit eukaryotic cells such as Un1Cas12f1, SpCas12f1 and AsCas12f1. The activity of our nucleases varies depending on the tracrRNA design as well as on the length of the repeat and of the spacer. μCas nucleases are a valuable addition to the CRISPR-Cas toolbox due to their expanded PAM recognition and miniature size. The extremely small size and high activity of the μCas nucleases give more flexibility in designing therapeutic AAV constructs. The additional space creates opportunities for more advanced regulatory and safety elements like anti-CRISPR proteins to further control off-target editing.

## Acknowledgments

We would like to thank Prof. Joseph Bondy-Denomy for critical reading of the article.

## Authors’ Contributions

A.S and L.A.T. contributed to the study design, experiments, data analysis, and article writing. Z.M and J.S-A. contributed to experiments and data analysis, T.C. and D.R. contributed to study design. M.S. contributed to study design, data analysis, and article writing.

